# A poly(A) isoform–aware single-cell and spatial atlas defines fibroblast niches in the human bladder

**DOI:** 10.64898/2025.12.18.695268

**Authors:** Briana A. Santo, Emily E. Fink, Pierre-Emmanuel Desprez, Yi-Chia Lin, Mohamed Eltemamy, Alvin Wee, Ninh B. Le, Alexandra Krylova, Uyen Tran, Veena Kochat, Kunal Rai, Douglas W. Strand, Oliver Wessely, Byron H. Lee, Angela H. Ting

## Abstract

Bladder function relies on coordinated interactions among epithelial, stromal, vascular, and neural compartments, but high-resolution molecular and spatial features remain undefined. We generated a publicly accessible, poly(A) isoform–aware single-nucleus and spatial reference of the adult human bladder spanning four anatomical regions and both sexes. Integrating 74,694 snRNA-seq profiles with 168,476 Xenium-resolved cells, we identified 22 cell types and 23,489 polyadenylation sites, with isoform usage improving stromal resolution beyond gene expression alone. Spatial mapping revealed layered fibroblast niches aligned with epithelial, vascular, and neural structures, supported by Visium data. This multimodal reference links isoform regulation to anatomical context and provides a reusable framework for cell-type annotation, cross-study integration, and analyses of bladder physiology and disease.

## Main Text

The bladder stores and expels urine through the coordinated function of a stratified urothelium, underlying stroma, and dense neurovascular network. Disruption of this organization contributes to common disorders, including interstitial cystitis and urothelial carcinoma (*1–3*). Loss of epithelial stratification, fibroblast remodeling, and altered innervation or vascular tone underlie symptoms such as urinary frequency, urgency, pain, and hematuria, and can drive fibrosis and loss of compliance in benign disease. Malignant transformation and tumor invasion arise through distinct processes, but the field lacks a unified, high-resolution molecular and spatial reference of the normal adult bladder from which to chart these trajectories (*4, 5*).

Recent single-cell studies have outlined major bladder cell types and provided initial views of urothelial stratification, fibroblast diversity, and immune composition in health and disease (*5–8*). However, two gaps remain: spatial context has not been resolved at single-cell resolution to define how cell types assemble into functional niches, and analyses have focused on gene-level counts while leaving alternative polyadenylation (APA) – a widespread post-transcriptional mechanism that influences mRNA stability, localization, and protein output – completely unexplored. In many tissues, APA can distinguish closely related cell states (*9–11*). However, its contribution to bladder cell identity, stromal heterogeneity, and tissue architecture has not been defined within an integrated spatial framework. A dataset that couples transcriptional profiles, isoform usage, and spatial organization is therefore needed as a reusable reference.

Here, we present a poly(A) isoform–aware single-nucleus and spatial atlas of the adult human bladder that integrates molecular, spatial, and post-transcriptional features at single-cell resolution across male and female donors and four anatomical regions: dome, neck, ureteral orifice, and ureterovesical junction (*12*). The atlas comprises 74,694 snRNA-seq nuclei and 168,476 spatially resolved cells, quantifying 23,489 poly(A) sites from 15,793 expressed genes, with 60.7% undergoing APA. Concordant cell-type and spatial patterns across snRNA-seq, Xenium, and Visium platforms support the robustness of this reference. Organized as a publicly accessible resource, the atlas provides standardized annotation and a foundation for downstream analyses of bladder physiology and disease.

This work contributes to the NIH Genitourinary Developmental Molecular Anatomy Project (GUDMAP) and supports GUDMAP’s goal of delivering high-resolution molecular and anatomical maps of the genitourinary tract (*13, 14*). By providing a poly(A) isoform-aware single-cell and spatial reference for the adult human bladder, this atlas establishes a framework to interrogate bladder organization, regulatory complexity, and disease mechanisms at unprecedented resolution. All data and code are publicly available through CELLxGENE Discover (*15*), GEO, SRA, and GitHub to facilitate cross-study comparison and future disease-focused investigations.

### Single-nucleus transcriptomic atlas of the adult human bladder

To construct a cellular atlas of the human bladder, we used single-nucleus RNA sequencing (snRNA-seq) to profile nuclei from four anatomical regions: the dome, neck, ureteral orifice (UO), and ureterovesical junction (UVJ). Histologically normal tissue was obtained from five organ donors and processed immediately after procurement. After quality control, 74,694 high-quality nuclei were retained across all regions (Fig. 1A; table S1). Unsupervised clustering identified 13 major groups, which were refined into 22 transcriptionally distinct cell types through compartment-level analysis guided by canonical markers (*16*) (Fig. 1B).

**Fig. 1.**
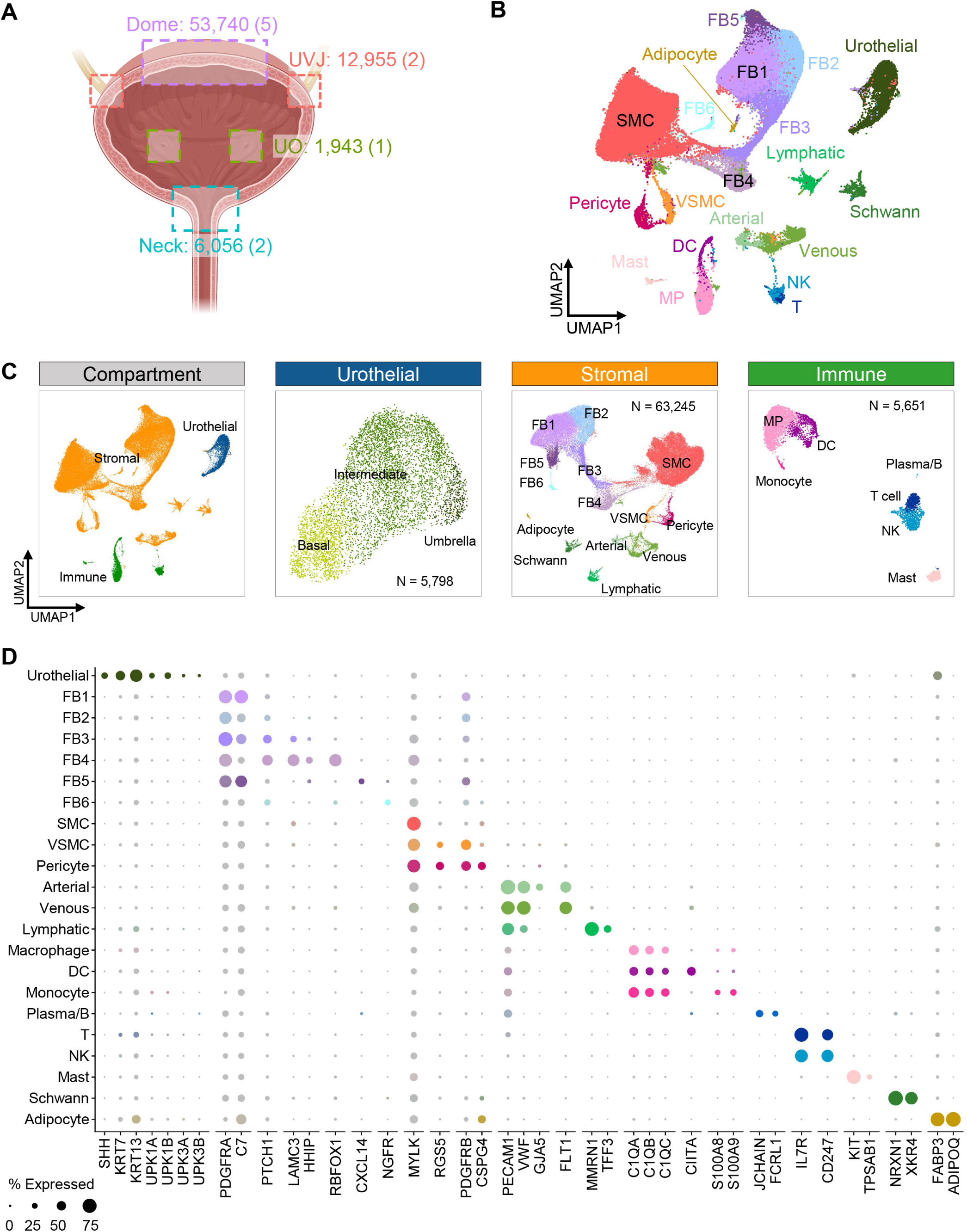
Cellular composition of the human bladder resolved by snRNA-seq. (**A**) Number of nuclei profiled per bladder region (dome, neck, ureteral orifice [UO], and ureterovesical junction [UVJ]). (**B**) Global UMAP embedding of 74,694 nuclei colored by 22 transcriptionally distinct cell types. (**C**) Compartment-level views highlight urothelial, stromal, and immune subsets. (**D**) Dotplot of canonical marker genes used for cell type annotation. Dot size indicates percent of nuclei expression the gene; color intensity reflects scaled expression. Dots are colored according to the corresponding cell type in the global UMAP. Abbreviations: SMC, smooth muscle cell; FB, fibroblast; VSMC, vascular smooth muscle cell; NK, natural killer cell; DC, dendritic cell; MP, macrophage.

The urothelium resolved into three layers corresponding to basal (*KRT5*^+^/*KRT13*^+^/*ADAMTS9*^+^), umbrella (*UPK1A*^+^/*UPK3A*^+^), and intermediate (*KRT13*^+^/*ABCC3*^+^) layers (Fig. 1C; fig. S1A; table S2). Stromal cells dominated the dataset (n = 63,245) and exhibited extensive heterogeneity, including six fibroblast subtypes distinguished by unique marker combinations as well as multiple smooth muscle and endothelial populations (Fig. 1C; fig. S1B). Smooth muscle cells segregated into general (*ACTA2^+^/MYLK^+^*), vascular (*RGS5^+^*), and pericyte (*CSPG4^+^*) subsets, while endothelial cells partitioned into arterial (*GJA5*^+^), venous (*FLT1*^+^), and lymphatic (*MMRN1*^+^/*TFF3*^+^) types. Additional stromal populations included Schwann cells (*NRXN*^+^/*XKR4*^+^) and adipocytes (*FABP3*^+^/*ADIPOQ*^+^). Immune cells, though less abundant (n = 5,651), encompassed macrophages, dendritic cells, monocytes, plasma/B cells, T cells, NK cells, and mast cells (Fig. 1C-D; fig. S1C). Collectively, these analyses define a comprehensive set of 22 transcriptionally distinct cell populations.

Regional comparisons revealed significant proportional differences (fig. S1D-F), including enrichment of smooth muscle cells in the bladder dome relative to the UVJ. Sex-specific analysis showed mast cells, urothelial cells, venous endothelium, arterial endothelium, and pericytes were more abundant in male samples, whereas macrophages, dendritic cells, lymphatic endothelial cells, and fibroblast subsets FB1 and FB3 were enriched in female samples (fig. S1G-I). These comparisons suggest anatomical and sex-associated variation in cellular composition.

### Polyadenylation isoform landscape across bladder cell types

Cell-state-specific gene expression is shaped not only by transcriptional control but also by post-transcriptional mechanisms that diversify the transcriptome. Among these, alternative cleavage and polyadenylation (APA) generates distinct mRNA isoforms that alter coding potential or 3’ untranslated region (UTR) length, thereby influencing protein sequence, mRNA stability, localization, and translational output (*17*). Approximately 80% of mammalian genes undergo APA, with polyadenylation [poly(A)] site choice regulated in tissue- and cell-specific manners to fine-tune gene expression (*18, 19*). APA has been linked to developmental transitions (*20–22*), immune activation (*23, 24*), and differentiation in multiple systems (*25–27*). However, its contribution to bladder cell identity has not been examined at cellular resolution, and no isoform-aware reference exists for this tissue. Capturing isoform-level variation alongside overall gene expression offers a more complete view of transcriptome architecture and adds a regulatory dimension to single-cell atlases.

To quantify poly(A) isoform expression, we applied scPASU (*28*), a single-cell workflow for transcriptome-wide APA analysis, to our snRNA-seq data (fig. S2A). We generated a harmonized poly(A) site reference across 20 bladder cell types (excluding plasma/B cells and monocytes due to low abundance), comprising 23,489 isoforms from 15,793 genes (table S3). Each cell type expressed distinct numbers of isoforms in addition to differences in total expressed genes, suggesting that isoform-level analysis provides additional resolution beyond gene counts alone (Fig. 2A).

**Fig. 2.**
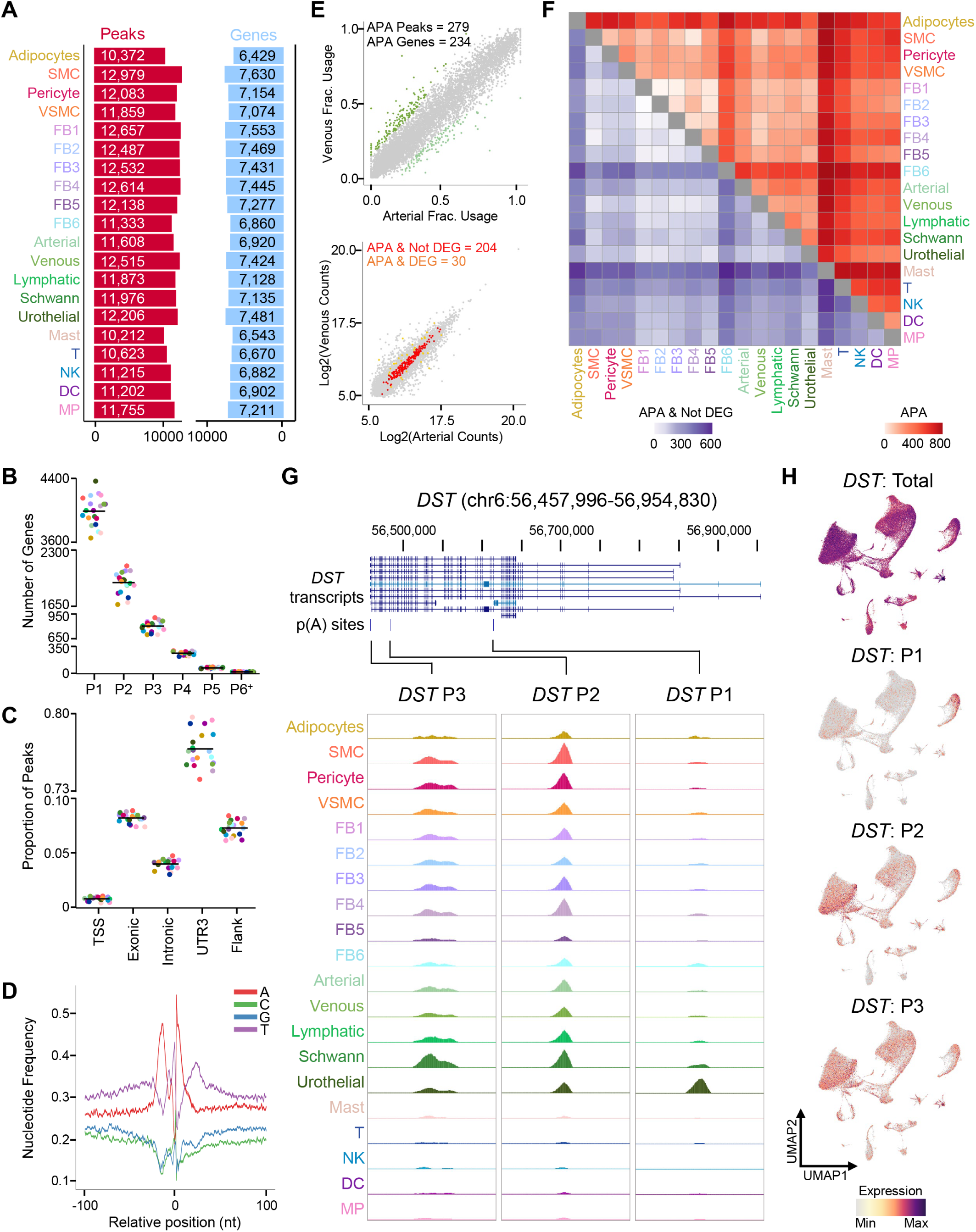
Poly(A) isoform quantification across bladder cell types. (**A**) Bar plot showing the number of distinct poly(A) isoforms (peaks) and expressed genes identified per bladder cell type.(**B**) Jitter plot showing the frequency of isoforms per gene across cell types. (**C**) Jitter plot showing the distribution of poly(A) sites across genomic contexts in each cell type. (**D**) Nucleotide frequency profiles within 100 bases upstream and downstream of poly(A) site start positions. (**E**) Scatter plots comparing arterial and venous endothelial cells: top panel shows 234 APA genes based on fractional usage; bottom panel highlights 193 APA genes without differential expression at the gene level (APA & Not DEG). (**F**) Heatmap summarizing APA gene counts (red) and APA-Not-DEG counts (purple) across all bladder cell types. (**G**) UCSC Genome Browser view of *DST* showing three poly(A) sites detected in snRNA-seq data. (**H**) Feature plots of *DST* total gene expression and isoform-specific expression projected onto the global snRNA-seq UMAP. Abbreviations: SMC, smooth muscle cell; FB, fibroblast; VSMC, vascular smooth muscle cell; NK, natural killer cell; DC, dendritic cell; MP, macrophage; APA, alternative polyadenylation; DEG, differentially expressed gene; Frac. Usage, fractional usage; TSS, transcription start site; UTR3, 3’ UTR.

Genes were divided into single- or multi- poly(A) genes, with the latter defined as those harboring two or more isoforms (Fig. 2B). Most poly(A) sites localized to 3’ UTRs (Fig. 2C), and flanking nucleotide profiles matched established distributions (*29*) (Fig. 2D). Poly(A) site positions along the gene length were broadly consistent across cell types (fig. S2B).

Differential isoform expression was assessed by computing fractional usage, defined as the proportion of reads mapping to a given isoform relative to all isoforms of the same gene within a cell type, and performing pairwise comparisons (table S4). For example, comparing arterial versus venous endothelium revealed 234 genes with isoform differences (Fig. 2E, top), including 204 genes where isoform variation occurred without changes in mRNA abundance (APA-non-DEG; Fig. 2E, bottom). Across all comparisons, APA differences were more frequent between distinct cell lineages than among closely related subtypes, such as arterial versus venous endothelial cells (Fig. 2F, red heatmap). These lineage-structured APA patterns persisted even when genes with differential expression were excluded (Fig. 2F, purple heatmap).

Examples of genes with cell type-specific isoform patterns include *DST*, *CD47*, and *DNAJB6*. *DST* exhibited three poly(A) isoforms (P1, P2, P3): P1 was enriched in urothelial cells, P2 across stromal populations with heightened expression in smooth muscle cells, and P3 in Schwann cells (Fig. 2G). When isoform counts were visualized on the original snRNA-seq UMAP, these distinct isoform-expression patterns were evident (Fig. 2H). *CD47* (fig. S2C) and *DNAJB6* (fig. S2D) each had two isoforms. *CD47*:P1 was enriched in urothelial cells and adipocytes, whereas *CD47*:P2 was broadly detected among stromal and urothelial populations. *DNAJB6*:P1 was observed across cell types with higher levels in adipocytes and urothelial cells, while *DNAJB6*:P2 was largely restricted to adipocytes.

### Spatial architecture of bladder cell types and microenvironments

To extend our molecular characterization of the bladder, we spatially mapped cell types and their interactions using Xenium In Situ. We designed a custom panel targeting both genes and isoforms and applied it to an independent bladder tissue sample (fig. S3A). Following quality control, 168,476 cells were retained, with mean gene/isoform and transcript counts of 80.3 and 183.8, respectively (table S5). Joint analysis of gene and isoform expression enabled clustering and cell type annotation, resolving 33 cell types (Fig. 3A; fig. S3B).

**Fig. 3.**
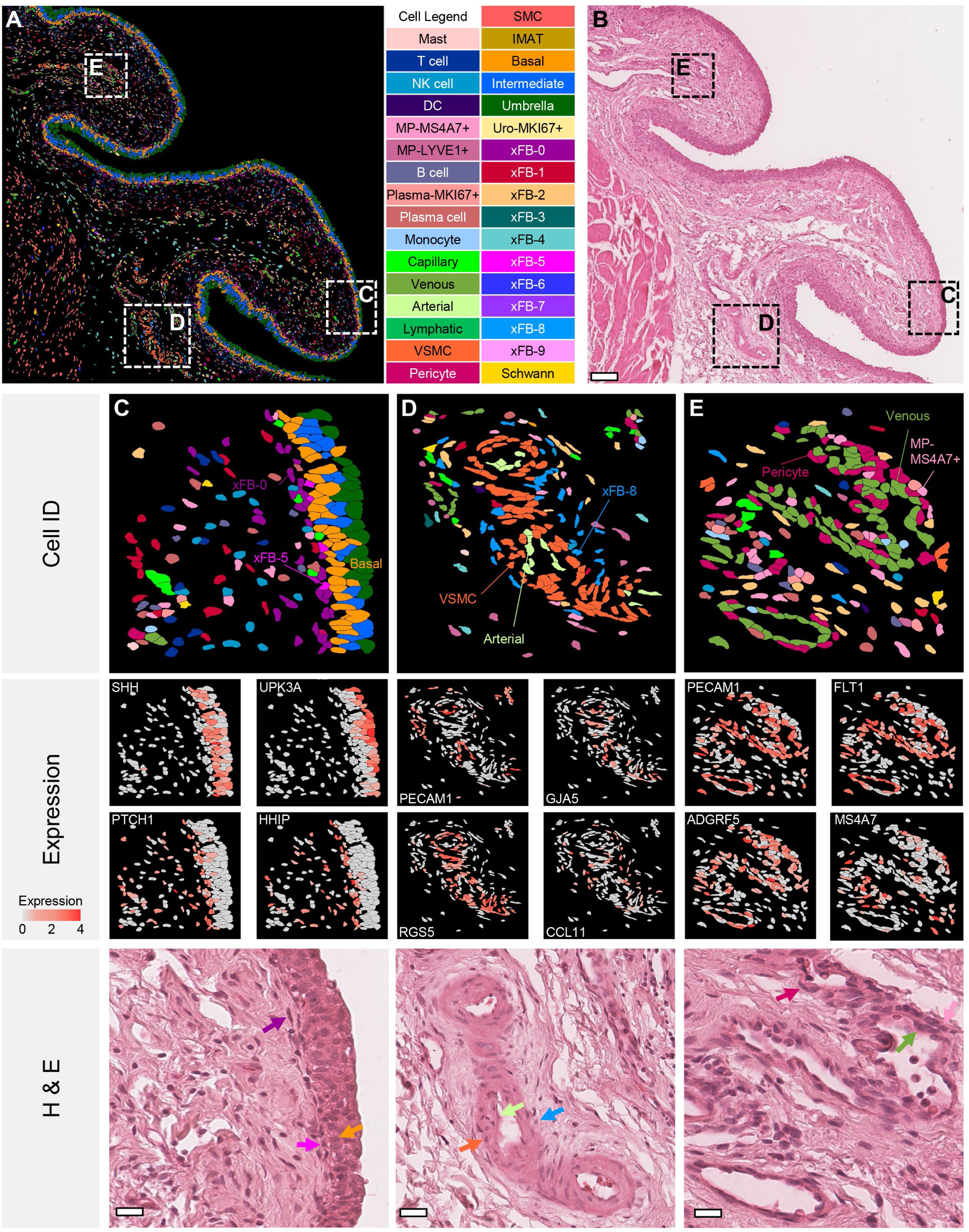
Spatial organization of bladder cell types in Xenium In Situ data. (**A**) Image dim plot of Xenium In Situ data showing all identified cell types across a bladder tissue section. Regions of interest (ROIs) highlighting microenvironments are outlined with white dashed boxes. (**B**) Corresponding H&E image of the same section; scale bar, 150 µm. (**C**) ROI at the urothelial-stromal interface showing basal urothelial cells (*SHH*), umbrella cells (*UPK3A*), and peri-urothelial fibroblasts xFB-0 (*PTCH1*) and xFB-5 (*HHIP*). (**D**) ROI featuring an artery with endothelial cells (*PECAM1*), arterial marker *GJA5*, vascular smooth muscle cells (*RGS5*), and xFB-8 (*CCL11*). (**E**) ROI featuring veins with endothelial cells (*PECAM1*), venous marker *FLT1*, pericytes (*ADGRF5*), and macrophages (*MS4A7*). For panels C-E, corresponding H&E images include color-coded arrows indicating cell types of interest; scale bars, 30 µm. Abbreviations: SMC, smooth muscle cell; xFB, Xenium fibroblast; VSMC, vascular smooth muscle cell; NK, natural killer cell; DC, dendritic cell; MP, macrophage; IMAT, intramuscular adipose tissue; Cell ID, cell type identity; H&E, hematoxylin and eosin.

Urothelial cells segregated into basal, intermediate, umbrella, and proliferative (*KI67*^+^) subpopulations. The stromal compartment revealed extensive heterogeneity, including 10 fibroblast subsets (xFB0-9), four endothelial cell types (arterial, venous, capillary, and lymphatic), three smooth muscle populations (SMC, VSMC, and pericytes), and single populations of intramuscular adipose tissue (IMAT) and Schwann cells. Immune cells encompassed B cells, plasma cells, *KI67*+ plasma cells, T cells, natural killer (NK) cells, dendritic cells (DC), monocytes, mast cells, and two macrophage subpopulations (*MS4A7*^+^ and *LYVE1*^+^). Differential expression analysis identified marker genes for each population (fig. S3C; table S6). To align molecular annotation with histologic context, we stained the tissue with H&E and registered the image to the Xenium fluorescence data for in-depth exploration (Fig. 3B).

We next examined microenvironments within the bladder, focusing on interfaces between parenchymal and stromal compartments and regions associated with vascular and neural structures. At the urothelial-stromal boundary (Fig. 3C), basal urothelial cells expressing sonic hedgehog (*SHH*) were juxtaposed with two fibroblast subsets: xFB-5 (*HHIP*^-^/*PTCH1*^+^) and xFB-0 (*HHIP*^+^/*PTCH1*^+^), consistent with known ligand-receptor distributions in this signaling pathway (*30*). In the lamina propria, the artery-associated regions (Fig. 3D) comprised arterial endothelial cells (*PECAM1*^+^/*GJA5*^+^), surrounding vascular smooth muscle cells (*RGS5*^+^), and fibroblast xFB-8 (*CCL11*^+^). Vein-associated regions (Fig. 3E; fig. S3D) included venous endothelial cells and neighboring capillaries (*PECAM1*^+^/*FLT1*^+^), surrounded by pericytes (*ADGRF5*^+^) and fibroblast xFB-9 (*CXCL14*^+^) as well as macrophages (*MS4A7*^+^). Regions surrounding innervating structures (fig. S3E) featured Schwann cells (*NRXN1*^+^) and a localized fibroblast subset xFB-7 (*DCN*^+^/*NGFR*^+^). Across these regions, spatial analysis revealed reproducible cellular arrangements in which stromal subsets aligned closely with epithelial, vascular, or neural structures.

### *In situ* validation of isoform-specific expression patterns

Having established the spatial organization of bladder cell types and stromal niches, we next leveraged the isoform-targeted Xenium panel to validate poly(A) isoform enrichment in situ, providing orthogonal confirmation of APA programs identified in snRNA-seq. This approach enabled us to examine whether isoform-specific expression patterns observed in transcriptomic data were preserved within tissue architecture.

We first focused on *DST*:P1, previously identified as highly specific to urothelial cells (Fig. 2G). Xenium analysis recapitulated this pattern, revealing strong enrichment of *DST*:P1 within the urothelial compartment and, more precisely, in basal cells, whereas total *DST* expression was uniform across all urothelial cell layers (Fig. 4A). Fractional usage analysis confirmed significant differences among basal, intermediate and umbrella cells, with *DST*:P1 fractional usage being the highest in basal cells (Fig. 4B), consistent with prior reports of *DST*:P1 localization in basal cells of the ureter (*28*). We next examined *MCCC2*:P1, another isoform associated with urothelial heterogeneity (fig. S4A). Similar to *DST*:P1, *MCCC2*:P1 exhibited preferential usage in basal cells (Fig. 4C). Joint density mapping of *DST*:P1 and *MCCC2*:P1 on the Xenium UMAP revealed near-exclusive co-localization within the basal cell cluster (Fig. 4D), indicating that isoform-level measurements add discriminatory power for closely related urothelial populations.

**Fig. 4.**
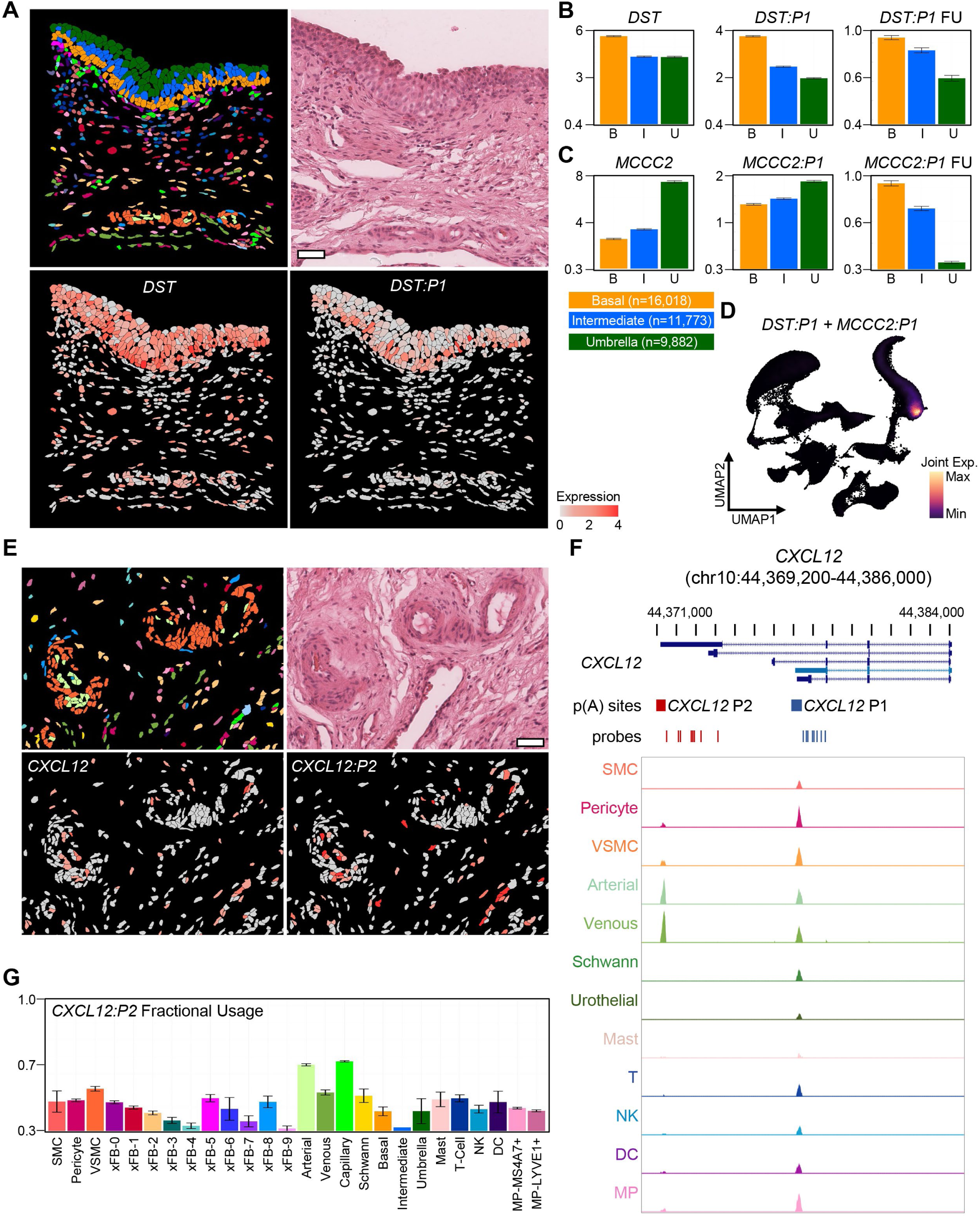
Spatial validation of poly(A) isoform expression in bladder microenvironments. (**A**) Xenium region of interest (ROI) with corresponding H&E image, feature plot of *DST* total genes expression, and *DST:P1* isoform expression; scale bar, 50 µm. (**B**) Bar plots showing DST total gene expression, *DST:P1* isoform expression and *DST:P1* fractional usage across urothelial subpopulations computed for the entire Xenium tissue section. (**C**) Bar plots showing *MCCC2* total gene expression, *MCCC2:P1* isoform expression, and *MCCC2:P1* fractional usage across urothelial subpopulations for the entire Xenium tissue section. (**D**) Joint density plot of *DST:P1* and *MCCC2:P1* isoforms projected onto the global Xenium UMAP. (**E**) ROI within the lamina propria centered on arteries and capillaries with corresponding H&E image, feature plots of *CXCL12* total gene expression, and *CXCL12:P2* isoform expression; scale bar, 45 µm. (**F**) UCSC genome browser view of *CXCL12* showing two poly(A) sites detected in snRNA-seq data across bladder cell types. (**G**) Bar plot of *CXCL12:P2* fractional usage across all cell types in the Xenium bladder tissue section. Abbreviations: SMC, smooth muscle cell; xFB, Xenium fibroblast; VSMC, vascular smooth muscle cell; NK, natural killer cell; DC, dendritic cell; MP, macrophage.

Isoform-level differences extended beyond the urothelium. *CXCL12*:P2, identified in snRNA-seq as enriched in arterial and venous endothelial cells, showed strong spatial specificity in Xenium data (Fig. 4E-F). Quantification across the entire tissue confirmed higher fractional usage of *CXCL12*:P2 in arterial and capillary endothelial cells (Fig. 4G). Notably, the apparent enrichment in venous endothelial cells observed in snRNA-seq likely reflected the presence of capillary endothelial cells within the venous cluster, a distinction clarified by spatial mapping. Feature plots further highlighted the restricted distribution of *CXCL12*:P2 to endothelial subsets, in contrast to the broad, non-specific expression observed at the gene level (fig. S4B-C). These spatial analyses demonstrate that isoform-restricted expression occurs in anatomically defined microenvironments and underscore the added resolution provided by isoform-aware profiling.

### Fibroblast architecture across the bladder wall

Building on the observation of distinct fibroblast subsets within bladder microenvironments, we next examined their spatial distributions across anatomical layers of the bladder wall. A region of interest spanning the bladder lumen through the urothelium and lamina propria to the upper muscularis propria was selected for high-resolution analysis (Fig. 5A-B). Within this region, six fibroblast subpopulations predominated: xFB-0 through xFB-5. When visualized sequentially by proximity to the bladder lumen, these populations revealed a highly ordered concentric arrangement, with concentric layers of fibroblast subsets increasing in depth from the urothelial surface (Fig. 5C). xFB-5 was positioned immediately adjacent to basal urothelial cells, followed by xFB-0. Deeper layers included xFB-1 in the upper lamina propria, xFB-2 in the mid-lamina propria, and xFB-4 in the outer lamina propria. In the muscularis propria, xFB-3 was enriched near the lumen-facing edge of the detrusor muscle, while xFB-6 localized around muscle fascicles.

**Fig. 5.**
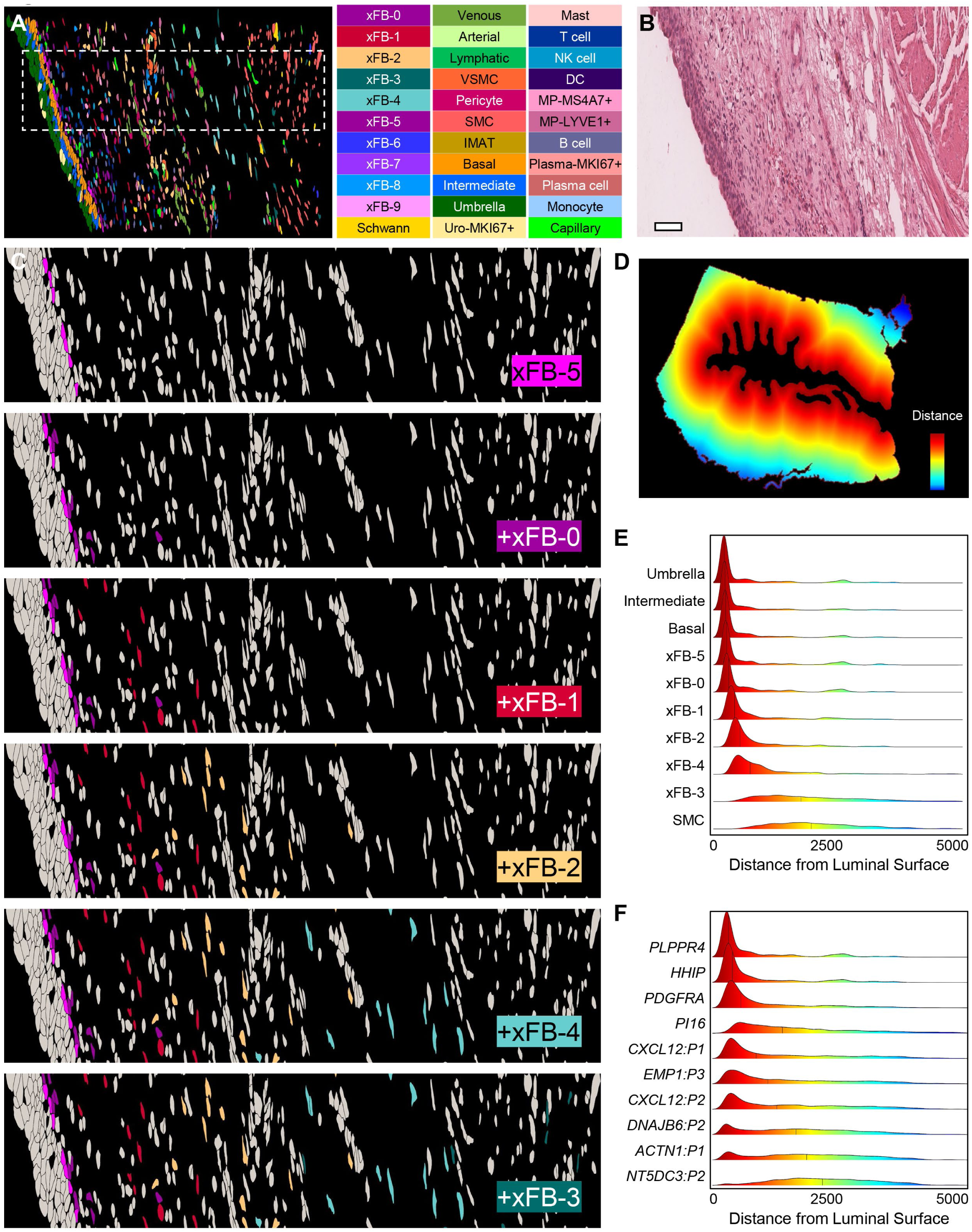
Spatial organization of fibroblast subpopulations in the human bladder. (**A**) A bladder region showing all identified cells. The region of interest (ROI) highlighted in panel **C** is outlined with a while dashed box. (**B**) Corresponding H&E image of the same bladder region in panel **A**; scale bar, 80 µm. (**C**) Organization of fibroblast subpopulations across anatomical layers: peri-urothelial region (xFB-5 and xFB-0), lamina propria (xFB-1, xFB-2, and xFB-4), and adjacent to the muscularis propria (xFB-3). (**D**) Heatmap showing pixel-wise distances from the bladder lumen across the entire tissue section, converted from pixels to microns. (**E**) Distribution of urothelial, fibroblast, and smooth muscle cell populations relative to luminal distance, color-coded as in panel **D**. (**F**) Expression profiles of generic and subpopulation-specific fibroblast marker transcrtips plotted relative to luminal distance, color-coded as in panel **D**. Abbreviations: SMC, smooth muscle cell; xFB, Xenium fibroblast; VSMC, vascular smooth muscle cell; NK, natural killer cell; DC, dendritic cell; MP, macrophage; IMAT, intramuscular adipose tissue.

To determine whether this layering was a localized or global feature, we quantified fibroblast distributions across the entire tissue section. A distance transform was computed for each pixel relative to the bladder lumen (Fig. 5D), and cell centroids were mapped to their corresponding luminal distances. This enabled visualization of cell type distributions as a function of depth (Fig. 5E). The concentric layering of fibroblast subpopulations was recapitulated at the tissue-wide scale, with xFB-5 and xFB-0 closely following basal urothelial cells, xFB-1, xFB-2, and XFB-4 spanning the lamina propria, and xFB-3 aligning with the detrusor muscle boundary.

Although prior studies have described general peri-urothelial, lamina propria, and intramuscular fibroblast types (*7, 31, 32*), the ability to distinguish multiple, spatially ordered fibroblast populations across these layers reflects the increased resolution afforded by combining gene- and isoform-level measurements. We evaluated whether the increased resolution may reflect the incorporation of isoform-level data into the Xenium panel and clustering workflow. To test this, we quantified the spatial distribution of differentially expressed genes and isoforms across fibroblast subsets using the same luminal distance framework. For each top marker gene and isoform, we computed transcript density relative to the bladder lumen and plotted feature distribution by cell type (Fig. 5F; fig. S5A).

Canonical markers such as *HHIP* aligned with peri-urothelial fibroblasts, while *PDGFRA* was broadly enriched across lamina propria subsets. In contrast, isoform-level features revealed sharper spatial distinctions. For example, *CXCL12*:P1 was enriched in both xFB-5 and xFB-3, marking fibroblasts adjacent to the urothelium and those bordering the detrusor muscle (Fig. 5F; fig. S5B; table S6). Meanwhile, *NQO1*:P2 showed selective enrichment in xFB-2, corresponding to the mid-lamina propria (fig. S5C). This combination of spatial mapping and isoform profiling reveals discrete fibroblast subpopulations arranged in a circumferential pattern across the bladder wall.

### Spatial neighborhood analysis of fibroblast niches

To further define cellular niches and refine fibroblast subpopulation identities, we performed spatial neighborhood analysis across the entire bladder tissue. Building on our observation of layered fibroblast organization and their proximity to epithelial, vascular, and neural structures, we quantified local cell-cell relationships to assess whether these spatial patterns were consistent across the tissue.

For each cell in the Xenium dataset, we computed pairwise distances to all other cells, identified its five nearest non-immune cell neighbors, and aggregated neighbor frequencies by cell type (Fig. 6A; table S7). This analysis revealed reproducible spatial associations. For example, basal urothelial cells were most frequently neighbored by other basal cells, followed by intermediate and umbrella cells, and xFB-0 and xFB-2. Similarly, xFB-1, localized to the upper lamina propria, was neighbored by xFB-0 and xFB-2, reflecting its transitional position between peri-urothelial and mid-lamina propria fibroblasts.

**Fig. 6.**
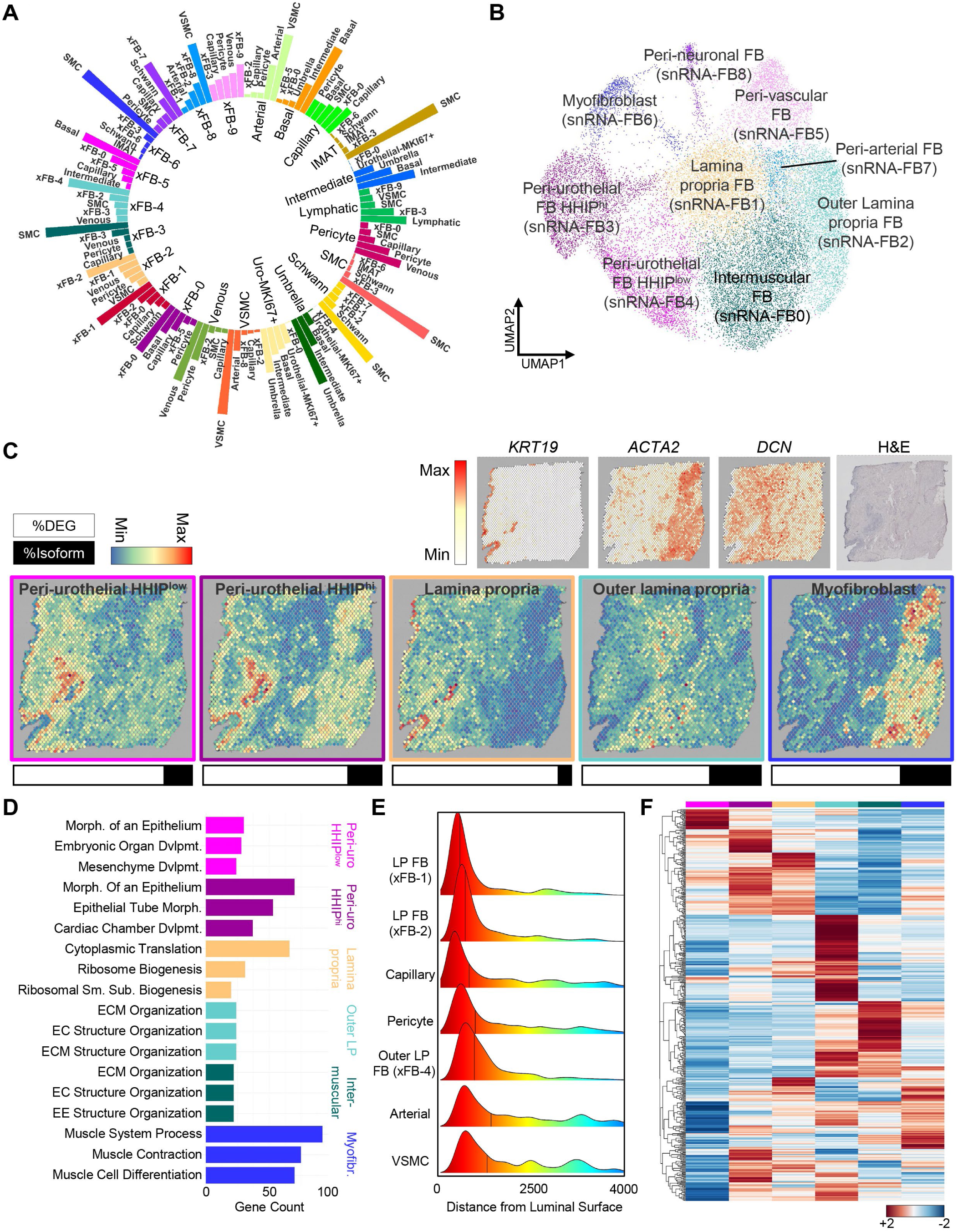
Multimodal integration resolves fibroblast niches and metabolic programs. (**A**) Circle plot summarizing nearest-neighbor analysis in Xenium data, showing the five closest non-immune neighbors for each cell type. (**B**) Multimodal clustering of snRNA-seq fibroblasts using combined gene expression and poly(A) isoform data. (**C**) Visium spatial transcriptomics of an independent bladder sample demonstrating reproducible fibroblast layering across peri-urothelial, lamina propria, and muscularis propria regions. Each panel shows combined expression of differentially expressed genes and isoforms for fibroblast populations identified in panel B, with bar plots below indicating relative contribution of genes and isoforms. Additional feature plots display canonical markers (*KRT19*, urothelium; *ACTA2*, muscle/myofibroblast; *DCN*, fibroblast) alongside the Visium H&E image. (**D**) Bar plot of top gene ontology (GO) terms for fibroblast layers based on snRNA-seq differentially expressed genes, arranged from peri-urothelial (HHIP^low^ and HHIP^hi^) to lamina propria, outer lamina propria, intermuscular, and myofibroblast populations. (**E**) Ridge plot showing Xenium spatial distribution of lamina propria fibroblasts and outer lamina propria fibroblasts relative to the bladder lumen. (**F**) Heatmap of metabolism-related genes per fibroblast population computed using the Recon3D database; normalized expression values were log2-transformed prior to plotting. Fibroblast layers are arranged as in panel **D**. Abbreviations: SMC, smooth muscle cell; xFB, Xenium fibroblast; snRNA-FB, multimodal fibroblast; VSMC, vascular smooth muscle cell; FB, fibroblast; %DEG, percentage differentially expressed gene; %Isoform, percentage isoform; Morph., morphogenesis; Dvlpmt., development; Sm., small; Sub., subunit; ECM, extracellular matrix; EC, extracellular; EE, external encapsulating; LP, lamina propria.

Beyond validating spatial proximity, neighborhood analysis supported an anatomical naming scheme for fibroblast subpopulations. We identified two peri-urothelial fibroblasts – xFB-5, directly adjacent to basal urothelial cells, and xFB-0, positioned immediately beneath – distinguished by *HHIP* expression level. Three lamina propria fibroblast subsets were resolved: xFB-1 (inner), xFB-2 (middle), and xFB-4 (outer). In the muscularis propria, xFB-3 and xFB-6 were localized around detrusor fascicles, consistent with intermuscular fibroblasts and myofibroblasts, respectively. Two vascular-associated fibroblasts were identified: xFB-8 (peri-arterial) and xFB-9 (peri-vascular). Finally, xFB-7 was enriched around Schwann cells, suggesting a peri-neuronal localization.

Recognizing that isoform-level data contributed to the resolution of these fibroblast subsets, we next asked whether re-clustering snRNA-seq data using both gene and isoform expression would yield similar granularity. We incorporated poly(A) isoform data into the original fibroblast Seurat object and performed multimodal clustering. This analysis resolved nine fibroblast subpopulations, closely matching those identified in Xenium data (Fig. 6B; table S8). To align clusters across platforms, we computed correlation matrices between normalized gene (fig. S6A) and isoform (fig. S6B) expression profiles from snRNA-seq and Xenium. This mapping enabled direct annotation of snRNA-seq clusters (Fig. 6B), with all Xenium-identified fibroblast types retained except for xFB-1 and xFB-2, which collapsed into a single lamina propria cluster (snRNA-FB1). This cross-platform alignment indicates that isoform-level measurements contribute substantially to resolving stromal heterogeneity.

To independently validate the spatial localization of these fibroblast populations, we analyzed an additional human bladder sample using the Visium spatial gene expression platform. Using enrichment scores derived from top differentially expressed genes and isoforms for each fibroblast subset, we recapitulated key features of fibroblast organization, including two peri-urothelial fibroblasts, two lamina propria subsets, and the myofibroblast population (Fig. 6C). These findings demonstrate that fibroblast layering is reproducible across platforms and specimens.

### Functional and metabolic specialization of fibroblast subsets

With the spatial localization of each fibroblast subset established, we next sought to functionally characterize these populations through gene ontology (GO) analysis of their differentially expressed gene sets (table S9). For the two peri-urothelial fibroblasts, enriched terms highlighted GO categories linked to epithelial-associated processes, including *Morphogenesis of an Epithelium* and *Mesenchyme Development* (Fig. 6D), consistent with their proximity to basal urothelial progenitor cells (Fig. 5C). Lamina propria fibroblasts exhibited distinct functional profiles: upper and middle subsets, which were consolidated as a single lamina propria fibroblast cluster, were enriched for transcriptional regulation (*Cytoplasmic Translation*), while outer lamina propria fibroblasts showed enrichment for extracellular matrix (ECM) remodeling (*ECM Structure Organization*), paralleling the ECM-related signatures observed in intermuscular fibroblasts (*ECM Organization*). Myofibroblasts, localized around detrusor muscle fibers, were uniquely enriched for terms related to smooth muscle biology, including *Muscle Cell Differentiation* and *Muscle Contraction*, reflecting their anatomical context.

Fibroblasts associated with innervating structures also displayed functional coherence with their spatial location. Peri-neuronal fibroblasts, surrounding Schwann cells (Fig. 3E), showed GO terms such as *Axonogenesis* and *Axon Guidance* (fig. S6C), suggesting a neuro-supportive role. Peri-vascular and peri-arterial fibroblasts were enriched for vascular maintenance terms, though their top annotations included more generic fibroblast functions such as ECM Organization and Response to Steroid Hormone (table S9; fig. S6C). Across fibroblast subsets, functional signatures aligned closely with anatomical position and neighboring cell types.

Given that the bladder’s vascular supply enters at the lamina propria, we hypothesized that the concentric layering of fibroblast subpopulations may reflect metabolic gradients shaped by vascular proximity. To test this, we plotted luminal distances of lamina propria fibroblasts alongside endothelial and VSMCs (Fig. 6E). Outer lamina propria fibroblasts (xFB-4) were closest to arterial endothelial cells, while upper (xFB-1) and middle (xFB-2) subsets aligned more closely with capillaries, indicating positional differences that align with distinct metabolic gene programs.

To further explore this, we clustered fibroblasts using a curated metabolism gene set from the Recon3D database (n = 2,248 genes). Each fibroblast subset was enriched for a distinct metabolic module (Fig. 6F), indicating distinct metabolic programs. To assess whether metabolic signatures alone could recapitulate fibroblast spatial identities, we computed enrichment scores using Recon3D genes and visualized them as Visium feature plots. While these profiles were less distinct than gene and isoform-based signatures, they nonetheless captured regional differences in fibroblast localization (fig. S6D), indicating that metabolic state varies systematically across fibroblast layers.

To quantify metabolic activity, we applied METAFlux, a computational tool for inferring metabolite flux from transcriptomic data (*33*). For each fibroblast subset, normalized flux values were computed to estimate relative uptake and efflux of metabolites (fig. S6E). Outer lamina propria fibroblasts exhibited the highest magnitude of flux values, indicating higher predicted flux activity in this stromal compartment. Together, these analyses indicate that fibroblast subpopulations across the bladder wall differ in functional and metabolic programs that align with their spatial positions.

## Discussion

This work delivers a unified, isoform-aware molecular and spatial reference for the normal adult human bladder, by integrating single-nucleus transcriptomics, spatial profiling, and isoform-resolved *in situ* analysis. Coupling gene-level expression with poly(A) isoform usage across complementary modalities provides a high-resolution framework that defines cell identity, tissue organization, and post-transcriptional regulation within a single resource. Concordant results across snRNA-seq, Xenium, and Visium underscore the robustness of this atlas. A central advance is the incorporation of isoform-level measurements, which increase cell-state resolution beyond gene counts and map consistently onto spatially defined niches. Together, these features establish a standardized baseline for cell-state annotation and comparative analyses of healthy and diseased bladder tissue.

The atlas reveals a structured stromal landscape composed of discrete fibroblast populations arranged in concentric layers from the urothelial surface through the lamina propria to the muscularis propria. These strata align with epithelial, vascular, and neural interfaces to form reproducible microenvironmental units. The resulting cellular and spatial maps offer an anatomical scaffold for interpreting bladder physiology and serve as a reference for developmental, inflammatory, fibrotic, and neoplastic states.

By making isoform-resolved single-cell and spatial datasets publicly accessible, this resource enables harmonized annotations, reproducible analyses, and cross-technology comparisons. It provides a foundation for mechanistic studies of epithelial–stromal communication, vascular remodeling, fibrosis, and tumor progression, and supports benchmarking new datasets, projection of disease states, and evaluation of isoform regulation across diverse pathologies. In sum, by establishing a high-resolution molecular and spatial baseline of the normal adult bladder, this atlas supplies the context needed to systematically map how benign remodeling and malignancy arise from normal tissue.

## Acknowledgments

We thank the organ donors and their families for gifting valuable tissues, Lifebanc for their partnership in supporting this research, and Dr. Eyal Gottlieb for advice on the metabolism analyses.

## Author contributions

Conceptualization: BAS, OW, BHL, AHT

Methodology: BAS, BHL, AHT

Investigation: BAS, EEF, PED, YCL, ME, AW, NBL, AK, UT, VK, DWS, BHL

Visualization: BAS, BHL, AHT

Funding acquisition: OW, BHL, AHT

Supervision: KR, OW, BHL, AHT

Writing – original draft: BAS, BHL, AHT

Writing – review & editing: EEF, PED, YCL, ME, AW, AK, UT, VK, KR, DWS, OW

## Data and materials availability

Single nuclei RNA-seq (snRNA-seq) and associated gene and peak count matrices have been deposited at GEO at GSE270224 and GSE270225. The processed snRNA-seq data are also available for interactive viewing at https://cellxgene.cziscience.com/e/fb5eeccf-3c2a-473c-b6e3-7a1dda18d42d.cxg/. The Xenium In Situ data have been deposited at GEO at GSE297459, and the Visium data under GSE307744. All original code has been deposited on GitHub (https://github.com/AngelaTingLab; https://doi.org/10.5281/zenodo.17889705). Any additional information required to reanalyze the data reported in this paper can be requested from the corresponding authors.

## Supplementary Materials for

### Materials and Methods

#### Human Bladder Procurement

Donor tissue was obtained through a research collaboration with Lifebanc, a non-profit organization that coordinates organ recovery for transplantation across more than 80 hospitals in Northeast Ohio. Under this agreement, Lifebanc screens potential donors for eligibility, including verification of donor designation in accordance with applicable state laws. With family consent, Lifebanc provides the lower urinary tract, including the bladder, for research purposes. For this study, bladders were procured from five donors, including three males and two females (mean age 25.2 years). A total of ten samples were excised from distinct bladder regions, including the dome (n=5), neck (n=2), ureteral orifice (n=1), and ureterovesical junction (n=2).

#### Single nucleus isolation

Dissected bladder regions were embedded in optimal cutting temperature (O.C.T.) compound, snap frozen on dry ice, and stored at −80°C until use. Nuclei isolation followed the Kidney Precision Medicine Consortium (KPMP) protocol.^4^ Briefly, ten 40 µm-thick cryosections were placed in 1 mL of ice-cold nuclear extraction buffer (20 mM Tris pH 8.0, 329 mM sucrose, 5 mM CaCl_2_, 3 mM MgAc_2_, 0.1 mM EDTA, 0.1 % Triton X-100 with 0.1 % RNase Inhibitor). The tissue mixture was pipetted up and down at least 20 times using a P1000 pipette with a trimmed tip to dissolve the O.C.T., followed by 10 additional mixing with an untrimmed p1000 tip. The suspension was transferred to a dounce homogenizer on ice and homogenized on ice with 5 strokes using pestle A and 20 strokes using pestle B, taking care to minimize bubble formation. The homogenate was incubated on ice for 10 minutes, then passed through a 40 µm FLOWMI™ cell strainer into a new 15 mL conical tube. The sample volume was brought up to 10 mL with ice-cold PBSE (1x PBS, 1mM EGTA) and centrifuged at 900 x g for 10 minutes at 4°C to pellet nuclei. Finally, the pellet was resuspended in ice-cold PBSE containing 1% BSA for manual counting.

#### Sample processing with 10X Genomics and cDNA library preparation

Single nucleus suspensions were processed using the 10X Genomics Chromium Single Cell 3’ Reagents Kit v3.1. We aimed to recover 10,000 nuclei per sample unless the total available nuclei were fewer; all samples had >5,000 nuclei captured for library preparation and sequencing. Following the manufacturer’s protocol, the nucleus suspension, the gel beads, and the emulsion oil were loaded onto a Chromium Single Cell Chip G and processed on the Chromium Controller. Immediately after droplet generation, samples were transferred to a PCR 8-tube strip (USA scientific for reverse transcription using SimpliAmp thermal cycler (Applied Biosystems). Following reverse transcription, cDNA was recovered using the recovery reagent provided by 10X Genomics, cleaned using Silane DynaBeads, and amplified for 11 cycles. The amplified cDNA was purified with SPRIselect beads (Beckman Coulter). To assess cDNA concentrations, a 1:10 dilution of each sample was analyzed on an Agilent Bioanalyzer High Sensitivity chip. Final cDNA libraries were constructed according to the Chromium Single Cell 3’ Reagent Kit v3.1 user guide.

#### snRNA-seq gene expression data analysis

Libraries were pooled and sequenced to a target depth of 50,000 read pairs per nucleus. De-multiplexed FASTQ files were processed with Cell Ranger (v7.1.0) with the count pipeline, mapping reads to the pre-built reference genome refdatagex-GRCh38-2020A and GTF from GENECODE v32 (GRCh28.p13). Alignment summary statistics for each sample are provided in Supplemental Table 1. Downstream analysis was performed in R using Seurat 4.3.0.1 and an established analysis pipeline (*34, 35*). The raw dataset contained 118,415 nuclei and was filtered for low-quality nuclei based on QC thresholds determined from distribution plots. Specifically, nuclei with >10% mitochondrial reads, gene counts <720 or >12,000, or total mapped read counts <2,000 were removed, leaving 74,694 high-quality nuclei for analysis. Seurat’s standard workflow was applied. Data were batch corrected using CCA and the IntegrateData function. Principal components (PCs) were computed from the 2,000 most variable genes, and the first 25 PCs were used for clustering at a resolution of 0.1. Clusters were visualized with UMAP and annotated based on marker gene expression. Cluster similarity was assessed using Peason correlation of the top 2,000 variable genes, grouping clusters into urothelial, stromal, and immune compartments. Clustering optimization was performed using clustree and chooseR (*36, 37*).

#### snRNA-seq gene expression subset analysis

For each subset analysis, nuclei from clusters assigned to a given compartment (immune, stromal, or urothelial) were processed using the same analysis pipeline described above, starting from raw gene counts. Additional filtering and iterative clustering were performed to remove cross-compartment contamination. Potentially contaminating clusters (e.g., suspected doublets) were removed ‘if the top100 differential markers for the cluster included markers exclusively expressed in another compartment (e.g. *MYLK* appearing among top100 markers in an immune cluster)’ AND ‘the contaminating marker was expressed in >10% of nuclei in that cluster’. Whe these criteria were met, the subset was re-clustered using the same workflow, excluding the contaminating cluster(s). After filtering, the final dataset included 5,651 immune cells, 63,245 stromal cells, and 5,798 urothelial cells. Each compartment was re-clustered using the following parameters: 45 PCs and a resolution of 0.7 for immune, 25 PCs and a resolution of 0.3 for stromal, and 25 PCs and a resolution of 0.1 for urothelial. Cell types within each compartment were annotated using canonical and established markers. Summaries of cell counts and differential expression analysis are provided in **Supplemental Table 1** and **Supplemental Table 2**, respectively. To assess differential cell type enrichment by bladder region and donor sex we completed proportion tests using *scProportionTest* with false discovery correction for multiple simultaneous tests (*38*). With respect to donor sex, we completed a bladder region agnostic analysis as well as a dome-specific analysis for which we had near balance by sex (3 male and 2 female donor domes).

#### Poly(A) isoform quantification from snRNA-seq data

1. *BAM pre-processing* Poly(A) isoform usage was quantified using scPASU, a computational pipeline for mapping cell-specific polyadenylation sites in single-cell/single-nuclei RNA-seq data (*28*). For previously aligned 10X Chromium 3’ samples (cellranger v7.1.0), PCR duplicates and ambiguously mapped reads were removed using the dedup module of umi_tools (v.1.1.4) and samtools (*39, 40*). Reads primed by genomically encoded A-rich sequences were filtered using polyAfilter (*41*). Processed BAM files were subset by cell barcodes for each cell type identified in snRNA-seq analysis for peak calling (Step 2). Subset BAM files were converted to FASTQ files for later poly(A) peak count-by-cell matrix generation (Step 4).
2. *Peak calling and Cell-specific peak references* For each cell type, peaks representing potential poly(A) sites were identified using MACS2 (*42*). Peaks with ≥ 3 poly(A) junction reads were assigned to transcription units (TUs). Ambiguous, broad, or low coverage peaks were removed. The poly(A) processing region (PR) for each peak was then inferred from the poly(A) junction reads. In parallel, we also repeated peak calling with just the poly(A) junction reads to rescue any low-coverage poly(A) sites that might have fallen below the noise threshold of MACS2. The two peak sets were then combined to produce a unified peak set for further filtering. The unified peak set was further filtered based on gene annotations, inferred PR width, presence of poly(A) signal, and estimated usage. Remaining peaks were re-annotated by genomic context to produce a dataset-specific poly(A) site reference.
3. *Harmonization of peak references across cell types* Cell-specific peak references were harmonized into a bladder poly(A) isoform peak reference using GenomicRanges in R (*43*). All cell-specific peak references were catenated and overlapping peaks across cell types were merged. Unique peaks per gene were enumerated and renumbered, and genes were classified as multi- or single- pA genes. The harmonized peak reference was saved for downstream steps.
4. *Poly(A) count-by-cell matrix generation* The FASTQ files from Step 1 were processed with Cell Ranger (v7.1.0) using the harmonized poly(A) site reference to produce the peak count-by-cell matrix for each sample and cell type.
5. *Identification of alternative poly(A) site usage* Peak counts per cell population were normalized by dividing the raw peak counts by the number of cells in that population. Isoforms were filtered at the gene level by filtering out isoforms with zero total gene expression (sum of normalized peaks). A global filter was applied to remove peaks with normalized count below the global median. Fractional usage of each isoform was computed as normalized peak count divided by the sum of peak counts for all isoforms of that gene. APA testing was performed between all pairs of bladder cell types, including only multi-pA genes. A peak was considered differentially used if it had an absolute log2 fold change in fractional usage ≥ 0.58 and an absolute fractional usage difference ≥0.15. Differentially expressed isoforms were visualized as scatter plots, color-coded by cell type according to the global UMAP from snRNA-seq.
6. *Identification of alternative poly(A) site usage in the absence of differential gene expression* To identify isoforms showing APA without differential gene expression (APA not DEG), pairwise pseudo-bulk DEG analysis was performed using DESeq2 (v.1.38.3) (*44*). For each comparison, three pseudo-replicates per group were generated by randomly sampling 70% of the nuclei without replacement. We used the Linear Ratio Test of DESeq2, where a Gamma-Poisson Generalized Linear Model was fitted and counts were normalized using the median ratio method. Benjamini-Hochberg correction was applied to p-values. Genes were considered differentially expressed if adjusted p-value <0.01 and absolute log2 fold change ≥ 0.58. Genes demonstrating APA in Step 5 and no differential expression in Step 6 were termed APA not DEG.

#### Characterization of poly(A) sites

Poly(A) sites in the reference were characterized using nucleotide frequency analysis, metagene profiling, and genomic context annotation. First, per-base nucleotide frequencies were calculated for the ±100 bp-region centered on the PR start coordinates in our poly(A) site reference and compared to those from the PolyASite atlas. Second, metagene plots were generated to show the distribution of poly(A) sites along TUs, where 0 represents the transcription start site (TSS) and 1 represents the transcription end site (TES), for both our reference and the PolyASite atlas. Lastly, peaks were annotated by genomic contexts in the following priority order to resolve ambiguous classifications: 3’ UTR, TSS-proximal, Exonic, Intronic, and Flank regions.

#### Xenium spatial gene expression assay and data analysis

##### Gene panel design

A custom Xenium human lower urinary tract probe panel (Panel ID: QK9QBG) targeting 354 genes and 125 mRNA isoforms was designed. The panel included markers for urothelial, fibroblast, immune, and smooth muscle cell markers as well as negative controls for cell type identification.

##### Xenium sample preparation

A 5 µm FFPE bladder sample from a deceased organ donor was sectioned onto a Xenium slide, fixed in 4% paraformaldehyde for 30 min, and rinsed in PBS. The tissue was permeabilized with 1% SDS for 2 min, washed twice in PBS, and immersed in pre-chilled 70% methanol for 60 min on ice, followed by two PBS washes. Custom probes and control probes (10nM) were hybridized overnight at 50°C in a thermocycler. After post-hybridization washes, probe ligation was performed at 37°C for 2 hr, followed by primer annealing. Rolling circle amplification was carried out for 2 hr at 37°C. Tissue section was then subjected to graded ethanol washes and incubated with blocking reagent to minimize non-specific antibody binding. Cell segmentation antibodies were applied to mark cell boundaries (ATP1A1 and E-cadherin), cell interior (18S rRNA, alpha SMA, and vimentin), and immune cells (CD45) and incubated overnight. Unbound antibodies were removed by washing, and staining intensity was enhanced using Xenium Staining Enhancer. After background fluorescence quenching and nuclear staining with DAPI, the slide was loaded onto the Xenium Analyzer.

##### Xenium Analyzer

Tissue sections were incubated with fluorescent-labeled probes and reagents provided by the manufacturer. The instrument performed 15 automated reagent cycles, imaging, and clearing between cycles. Z-stack images were acquired across the tissue thickness at 0.75-μm step size.

##### Image processing and analysis

Z-stack images were stitched using DAPI to create a composite Region of Interest (ROI) image for spatial mapping. Xenium Onboard Analysis detected transcripts as ‘puncta’, which is defined as observed photons in every cycle, image, and color. Emission profiles were fitted into a Gaussian distribution to generate unique optical gene signatures over 15 Xenium cycles. For each decoded transcript, a Q score was assigned, and transcripts with Q ≥ 20 were considered for Xenium spatial gene plots. Cell segmentation was performed using the Xenium Cell Segmentation Kit (10x Genomics) to generate a feature-cell matrix for downstream analysis. The segmentation algorithm integrates 3D DAPI information across all Z-slices, and optionally 2D focus images in the multimodal workflow, to improve accuracy. A flattened 2D segmentation mask was generated, with transcripts assigned to cells based on their X–Y coordinates. Nucleus and cell boundary polygons were provided as approximations of the segmentation masks for visualization in Xenium Explorer and other analysis software, and corresponding data were stored in the cells.zarr.zip output file.

##### Post-Xenium histology and image registration

H&E staining was performed on the same tissue sections. Whole-slide bright-field images were captured at 20x magnification using a Keyence (Itasca, IL) microscope and stitched together.

The H&E image was registered to the Xenium fluorescence image using the Xenium Explorer v3 [10X Genomics]. Briefly, the H&E image was converted to ome.tif format in python using the tifffile and opencv libraries, imported into Xenium Explorer, and aligned using landmark-based registration (*45, 46*).

##### Data QC

Raw Xenium bladder data were filtered for low-quality cells using thresholds of ≥5 detected genes and ≥80 transcripts per cell. Thresholds were determined by visualizing gene and transcript count distributions as histograms. The final dataset included 168,476 high quality cells, with median values of 134 transcripts and 69 genes (or isoforms) per cell, for downstream analysis.

##### Cell clustering and identification

Analysis was performed in R (v4.3.0) using Seurat (v5.1.0) with the standard workflow for image-based spatial data. Xenium data were normalized, PCs computed using all genes and isoforms, and clustering parameters optimized using clustree and elbow plots (*36*). Using 33 PCs and 0.4 resolution, 18 clusters were visualized with UMAP and annotated into four compartments, urothelium, stromal (non-fibroblast), fibroblast, and immune, based on marker gene expression. Marker genes included *KRT5* and *UPK3A* for urothelial cells, *ACTA2*, *MYLK*, *PECAM1*, *XKR4*, and *RGS5* for stromal cells, *PDGFRA* and *DCN* for fibroblasts, and *PTPRC* for immune cells. Subset analysis for each compartment followed the same workflow, starting from normalization and clustering optimization. Optimal parameters were:

- Urothelial: 20 PCs, resolution = 0.5
- Stromal: 14 PCs, resolution = 0.4
- Fibroblast: 11 PCs, resolution = 0.3
- Immune: 13 PCs, resolution = 0.3

Final clusters identified:

- 4 urothelial clusters (37,919 total cells)
- 9 stromal clusters (72,458 total cells)
- 10 fibroblast clusters (27,351 total cells)
- 10 immune clusters (30,748 total cells)

Cluster identities were confirmed by visualizing marker gene expression with violin and feature plots and computing differentially expressed genes and isoforms using *FindAllMarkers* in Seurat.

##### Computation of cell, gene, and isoform distances from the bladder lumen

The registered H&E image was loaded into MATLAB (R2024b), and a global mean-based threshold was applied to segment the bladder tissue and lumen regions from background. Luminal boundaries were identified using *bwboundaries*. The distance transform of the bladder tissue region was computed relative to the luminal edge using the *bwdist* function, where minimal values corresponded to surface urothelium and maximal values to the muscularis propria. Cell type distributions were calculated by extracting cell segmentation centroids from the Xenium object and computing their nearest luminal boundary point using *pdist2*. Distance was converted to microns using image resolution (microns/pixel). Cell type distributions were visualized as ridge plots. Gene and isoform transcript distributions were computed similarly using transcript centroids.

##### Isoform fractional usage

Fractional usage of APA isoforms on the Xenium panel was computed per cell type. For isoforms classified as 3’ UTR or Flank (e.g., *CXCL12:P1*), fractional usage equals normalized isoform expression divided by normalized total gene expression. For isoforms classified as TSS-proximal or intronic (e.g., *ITGA5:P1*), fractional usage equals normalized isoform expression divided by the sum of normalized isoform expression and normalized total gene expression. Mean and standard error across cells were calculated for cell population.

##### Xenium nearest-neighbor analysis

Nearest-neighbor analysis was performed using cell segmentation centroids extracted from the Xenium object. Pairwise distances between all cells were computed, and for each cell, the five nearest neighbors and their cell-type identities were recorded. Frequencies of neighbor identifies were aggregated across all cells of each population. This procedure was repeated for all cell types, and results were visualized using a circle plot.

### Multimodal clustering of snRNA-seq data incorporating poly(A) isoforms

Fibroblast snRNA-seq data were performed in R using the Seurat Weighted Nearest Neighbor Analysis (WNN) workflow. The gene expression matrix and scPASU-derived poly(A) isoform matrix for fibroblasts were combined into a single Seurat object. Each assay was pre-processed independently, including normalization (*NormalizeData*), variable feature selection (*FindVariableFeatures*), scaling (*ScaleData*), and dimensionality reduction (*RunPCA*).

Multimodal neighbors were computed using *FindMultimodalNeighbors*, which integrates RNA and poly(A) isoform similarities in a weighted manner. Clustering parameters were optimized to maintain consistency with the original snRNA-seq analysis. Final multimodal clustering used 15 PCs for gene expression, 25 PCs for poly(A) isoforms, and a resolution of 0.3, resulting in nine fibroblast subpopulations.

### Functional characterization of bladder fibroblast subsets

Differentially expressed genes were computed for each multimodal fibroblast subpopulation using the normalized RNA and poly(A) isoform assays. Features with log2FC > 0.58 and q-value < 0.05 were retained. Functional characterization included:

1. *Gene set enrichment analysis* Gene ontology analysis was performed using clusterProfiler to identify enriched biological processes for each fibroblast population (*47*). Processes were ranked by gene counts and q-values, and top terms were visualized as bar plots.
2. *Metabolic profiling using Recon3D* Metabolic differences were assessed using the Recon3D database, which consists of a curated list of 2,248 metabolism-related genes (*48*). The normalized RNA assay was subset to these genes, and differential expression analysis was performed using Seurat’s *FindAllMarkers* function. Genes with log2FC > 0.58 and adjusted p-value < 0.05 were considered significant. Metabolic heterogeneity was visualized by log2-transforming the subset RNA assay and applying hierarchical clustering with pheatmap package (*49*).
3. *Cell metabolic flux inference* METAFlux was applied to infer cell-specific metabolic flux from snRNA-seq data (*50*). Using the normalized RNA expression assay, “human_gem” database, 50 bootstraps, and a seed = 1, an average expression profile for stratified bootstrapped samples was created using the *calculate_avg_exp* function. Reaction activity scores and metabolic flux were calculated using the *calculate_reaction_score and compute_sc_flux* functions, respectively. Flux values were normalized, and “nutrient_lookup_files” were used to plot flux and exchanges for >60 metabolites and reactions (e.g., glucose uptake). Normalized flux matrices were averaged across bootstraps, reformatted to metabolite x cell type, and then used to calculate pathway level activity per cell type. Results were visualized as a heatmap.

### Visium spatial gene expression assay and data analysis

#### Visium spatial gene expression library generation

Bladder samples were collected immediately upon surgical removal, embedded in Optimal Cutting Temperature (OCT) compound, and stored in −80°C until ready to process (*51*). Following the Visium Spatial Tissue Optimization User Guide (CG000238 Rev D), an optimal permeabilization time of 6 minutes was determined for bladder tissue. Sections of 10 μm thick were placed on a Visium Spatial Gene Expression slide. Tissue fixation and H&E staining were performed according to the 10X Genomics Tissue Preparation Guide. Bright field images were taken with a Helios Slide Scanning Microscope (Leica Microsystems) at 10x magnification. Libraries were prepared per the Visium Spatial Gene Expression User Guide with the following parameters: 6min permeabilization time, 16 cycles for cDNA amplification, and 16 cycles for the sample index PCR. Libraries were pooled and sequenced to a target depth of 50,000 read pairs per tissue covered spot on the Capture Area.

#### Data processing and analysis

FASTQ files were processed using Space Ranger (v4.1.0). The count pipeline was executed twice using the pre-built GRCh38-2020A reference genome to generate a spot x gene matrix as well as using the scPASU harmonized reference to generate a spot x isoform matrix. Space Ranger was configured with the “reorient-images” option for optimal fiducial alignment. Aligned data was then processed in R (v4.3.0) using Seurat (v5.1.0). Gene and isoform assays were combined into a single Seurat object and normalized using *SCTransform*.

#### Spatial validation of fibroblast metabolic profiles

To validate distinct metabolic profiles of fibroblast populations, module scores were computed for multimodal fibroblast differentially expressed genes (DEGs), isoforms, and Recond3D-derived DEGs. Module scores were calculated per spot as follows. For each marker set, the Visium assay was subset to the corresponding expression submatrix. Within each submatrix, the top 15% of spots with the highest expression for each gene or isoform were retained, while all other values were set to zero. These filtered expression values were then aggregated, and per-spot averages across all genes/isoforms in the module were computed to generate a module score vector (one score per spot). This approach yielded a composite signature score enriched for the most highly expressing locations. Module scores were visualized across tissue sections using SpatialFeaturePlot.

---

**fig. S1.**
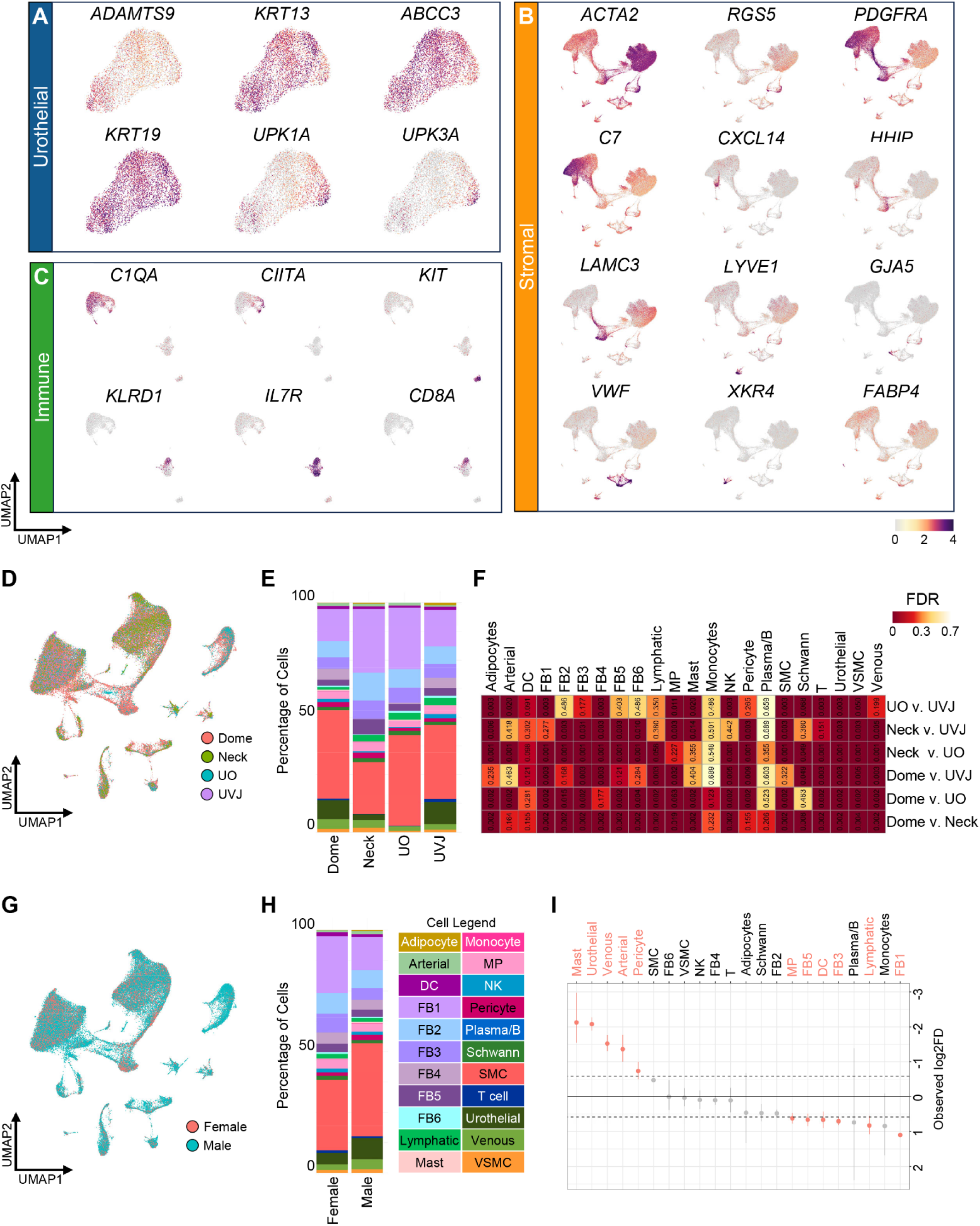
Compartment-level and regional analysis of bladder cell types by snRNA-seq. (**A**) UMAP of the urothelial compartment showing feature plots for canonical markers of basal, intermediate, and umbrella cells. (**B**) UMAP of the stromal compartment showing feature plots for markers for smooth muscle cells, vascular smooth muscle cells, fibroblasts, lymphatic endothelial cells, arterial and venous endothelial cells, Schwann cells, and adipocytes. (**C**) UMAP of the immune compartment showing expression of markers for macrophages, dendritic cells, mast cells, natural killer cells, and T cells. (**D**) Global UMAP colored by anatomical bladder region (UO, UVJ, neck, dome). (**E**) Bar plot summarizing cell type distributions across donor bladder regions. (**F**) Heatmap showing p-values (numbers) and false discovery rates (FDR; color) from cell type enrichment testing between bladder regions. (**G**) Global UMAP colored by donor sex. (**H**) Bar plot summarizing cell type distributions by sex. (**I**) Permutation plot showing relative enrichment of cell types between male and female donors. Positive log_2_ fold difference (Observed log2FD) indicates enrichment in female samples. Abbreviations: UO, ureteral orifices; UVJ, ureterovesical junction; SMC, smooth muscle cell; FB, fibroblast; VSMC, vascular smooth muscle cell; NK, natural killer cell; DC, dendritic cell; MP, macrophage.

**Fig. S2.**
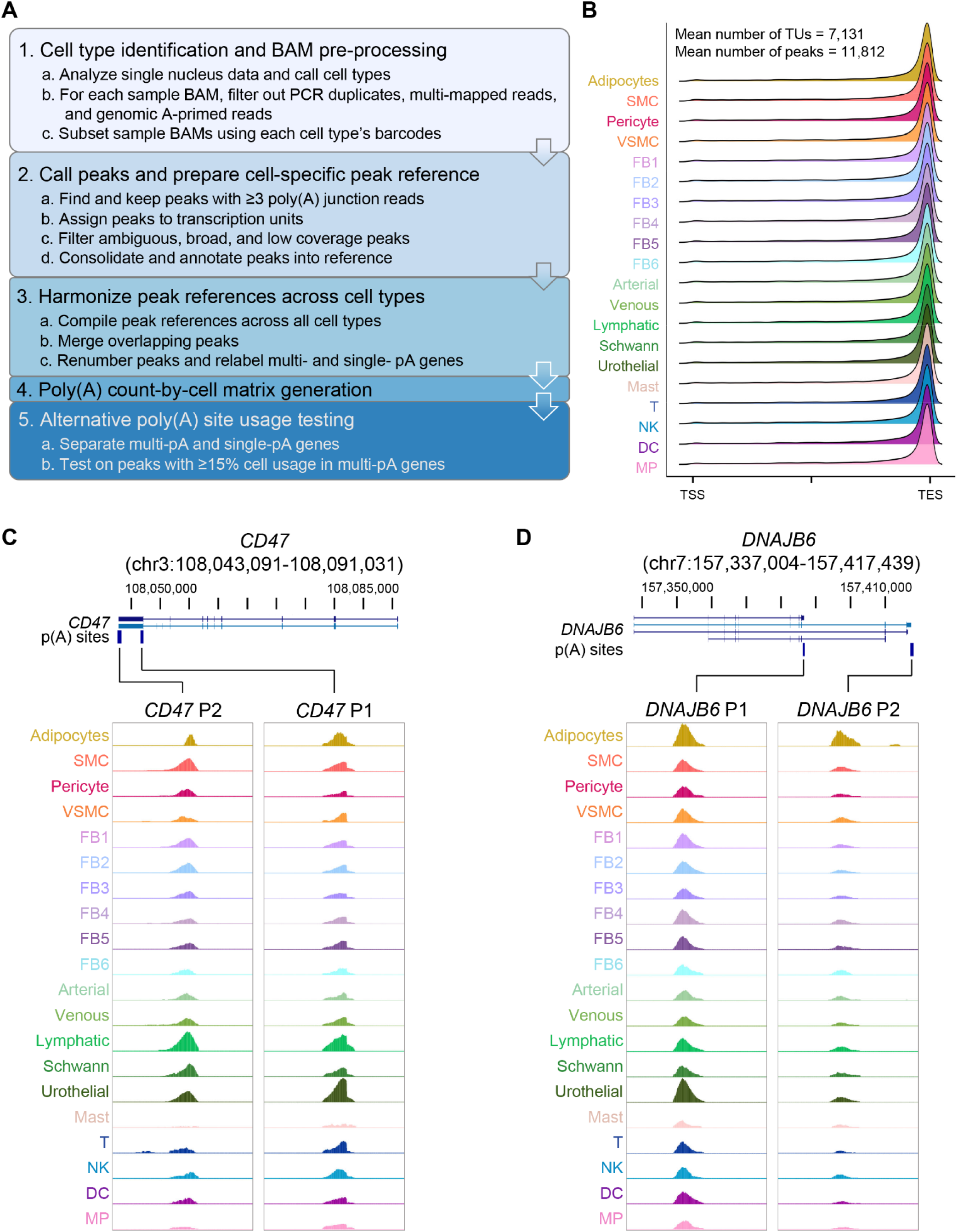
Workflow for poly(A) isoform quantification and representative bladder examples. (**A**) Schematic of the scPASU pipeline for single-cell polyadenylation site usage analysis, including harmonization of isoform references across all cell types. (**B**) Metagene plot showing the distribution of poly(A) sies across transcription units (TUs) for all cell types. (**C**) UCSC Genome Browser view of the CD47 locus showing tow distinct poly(A) sites detected in snRNA-seq data across bladder cell types. (**D**) UCSC Genome Browser view of the DNAJB6 locus showing two distinct poly(A) sites detected in snRNA-seq data across bladder cell types. Abbreviations: UO, ureteral orifices; UVJ, ureterovesical junction; SMC, smooth muscle cell; FB, fibroblast; VSMC, vascular smooth muscle cell; NK, natural killer cell; DC, dendritic cell; MP, macrophages.

**Fig. S3.**
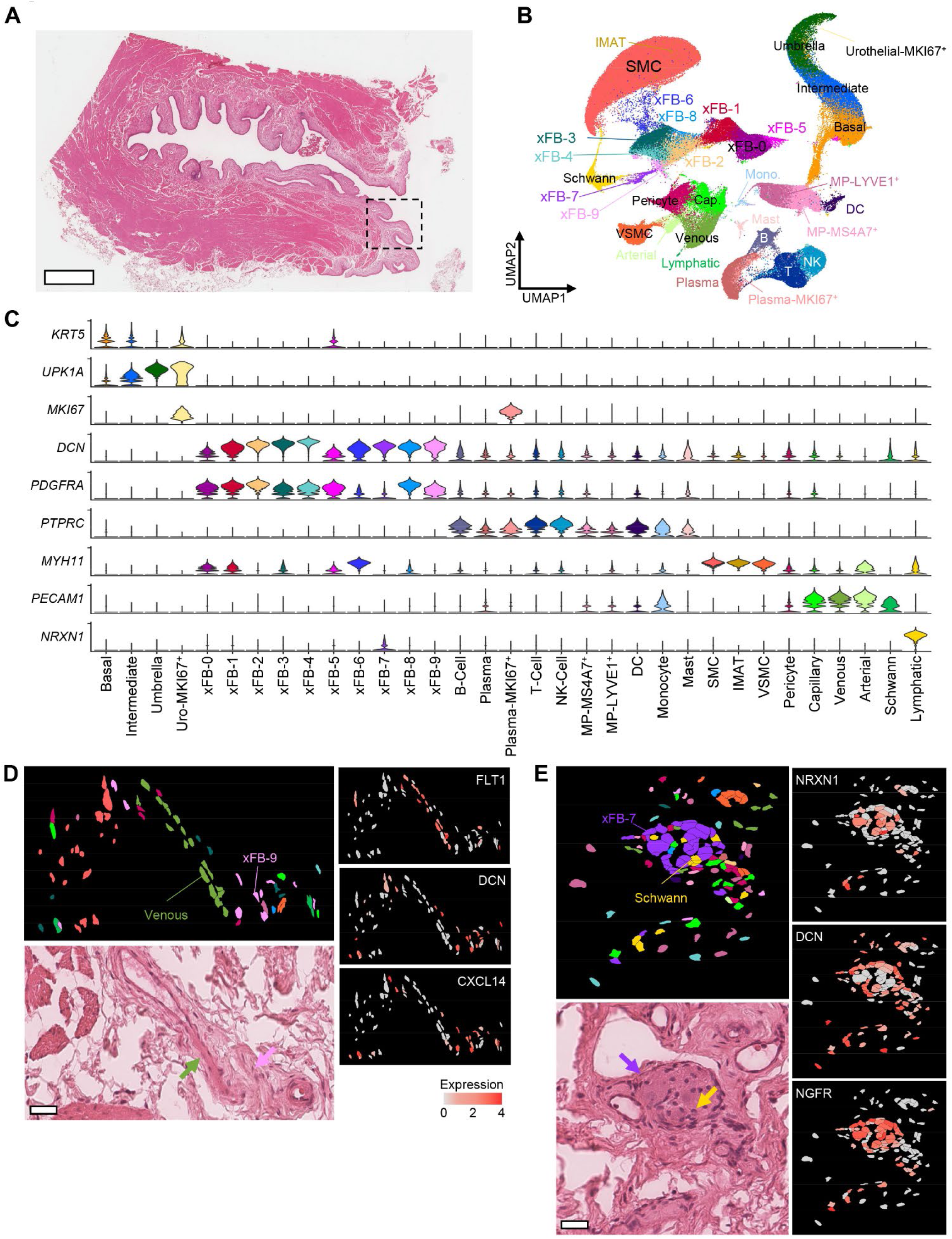
Global clustering and cell type identification in bladder tissue using Xenium In Situ data. (**A**) Whole-slide image of bladder tissue. Scale bar, 2 mm. Dashed box indicates the region shown in Fig. 3A. (**B**) Global UMAP of clustered Xenium data annotated with identified bladder cell types. (**C**) Violin plots showing expression of key marker genes used for cell type identification. Colors correspond to cell types shown in the global UMAP. (**D**) Region of interest (ROI) featuring a vein, with spatial expression of venous marker (*FLT1*), generic fibroblast marker (*DCN*) and xFB-9 specific marker (*CXCL14*). Scale bar, 30 µm. (**E**) ROI featuring a nerve bundle, with expression of nerve marker gene (*NRXN1*), generic fibroblast marker (*DCN*), and xFB-7 specific marker (*NGFR*). Corresponding H&E images are shown for panels D and E; scale bar, 30 µm. Abbreviations: SMC, smooth muscle cell; xFB, Xenium fibroblast; VSMC, vascular smooth muscle cell; NK, natural killer cell; DC, dendritic cell; IMAT, intramuscular adipose tissue.

**Fig. S4.**
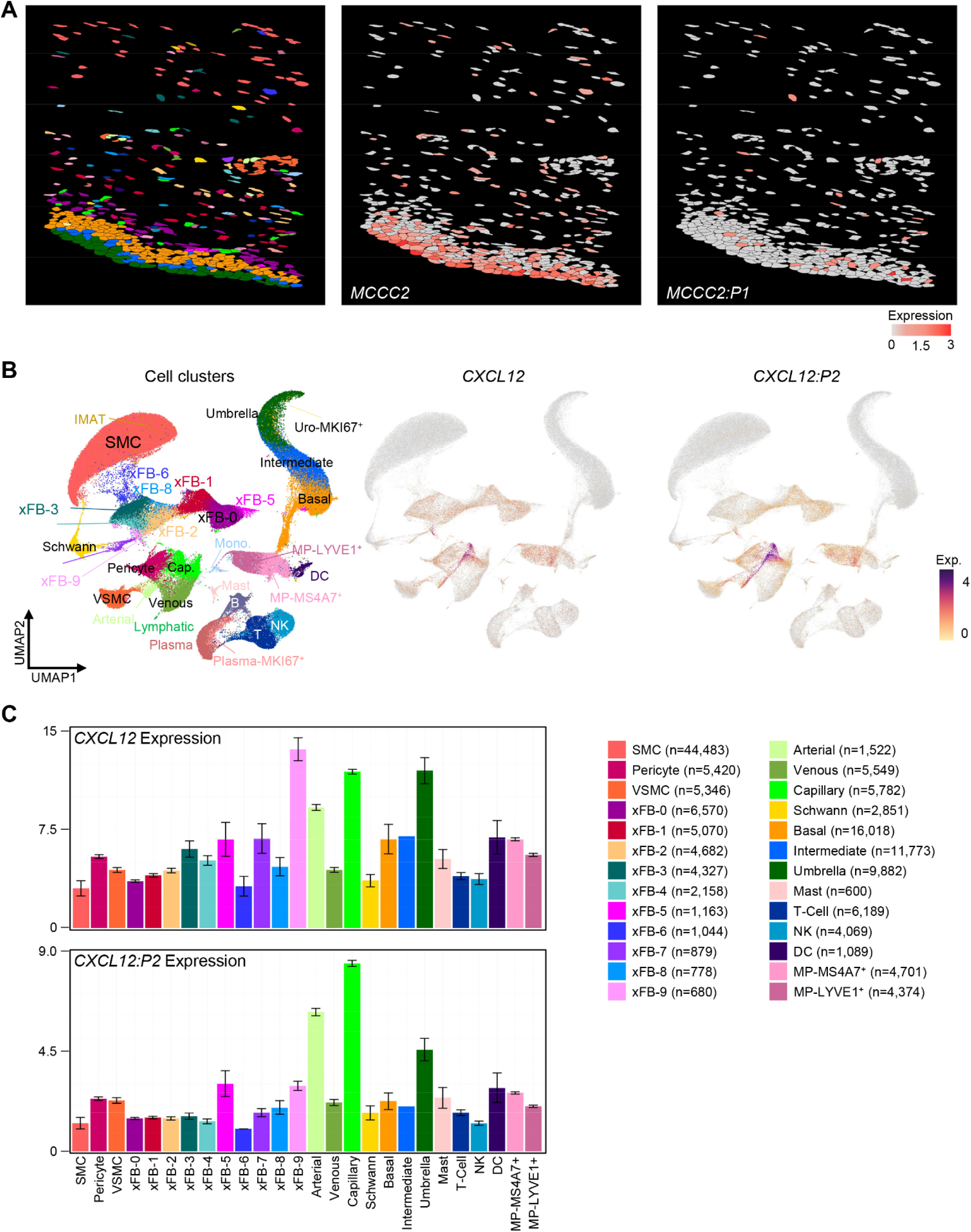
Poly(A) isoform usage across bladder cell types in Xenium In Situ data. (**A**) Region of interest (ROI) within the Xenium dataset focused on the urothelial compartment, showing total gene expression of *MCCC2* and spatial localization of the *MCCC2:P1* isoform. (**B**) Global UMAP displaying total gene expression of *CXCL12* and *CXCL12:P2* isoform expression. (**C**) Bar plots summarizing total *CXCL12* gene and *CXCL12:P2* isoform expressions across annotated cell types in the Xenium tissue. Abbreviations: SMC, smooth muscle cell; xFB, Xenium fibroblast; VSMC, vascular smooth muscle cell; NK, natural killer cell; DC, dendritic cell; IMAT, intramuscular adipose tissue.

**Fig. S5.**
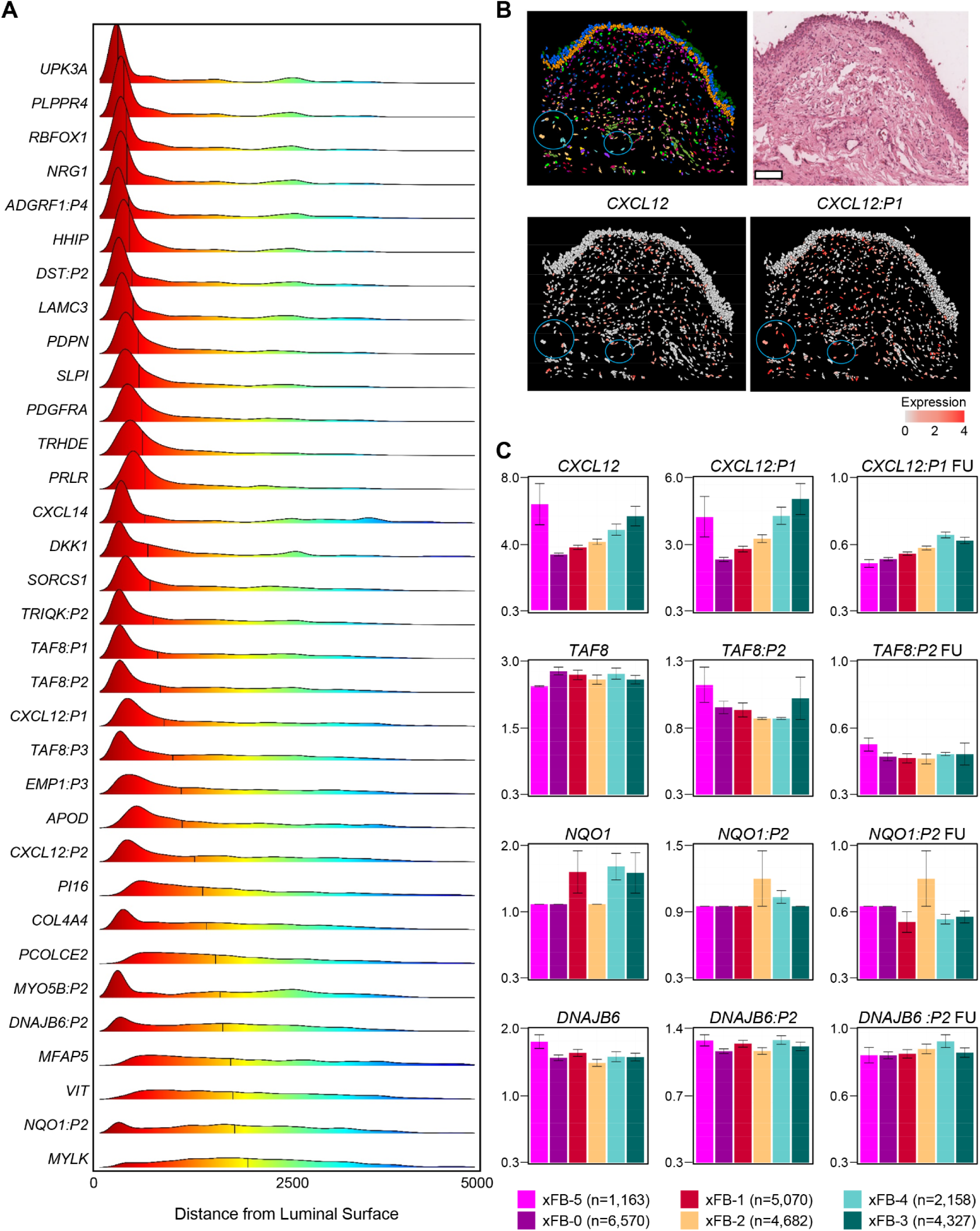
Alternative polyadenylation (APA) differences among fibroblast subpopulations across the bladder wall. (**A**) Distribution of fibroblast marker genes and differentially expressed poly(A) isoforms, along with their corresponding total gene transcripts, across spatially distinct fibroblast subpopulations. (**B**) Xenium region of interest with matched H&E staining, total gene expression of *CXCL12*, and spatial localization of the *CXCL12:P1* isoform. Scale bar, 100 µm. (**C**) Bar plots showing total gene and select isoform expressions and isoform fractional usage (FU) across fibroblast subpopulations. Abbreviations: xFB, Xenium fibroblast.

**Fig. S6.**
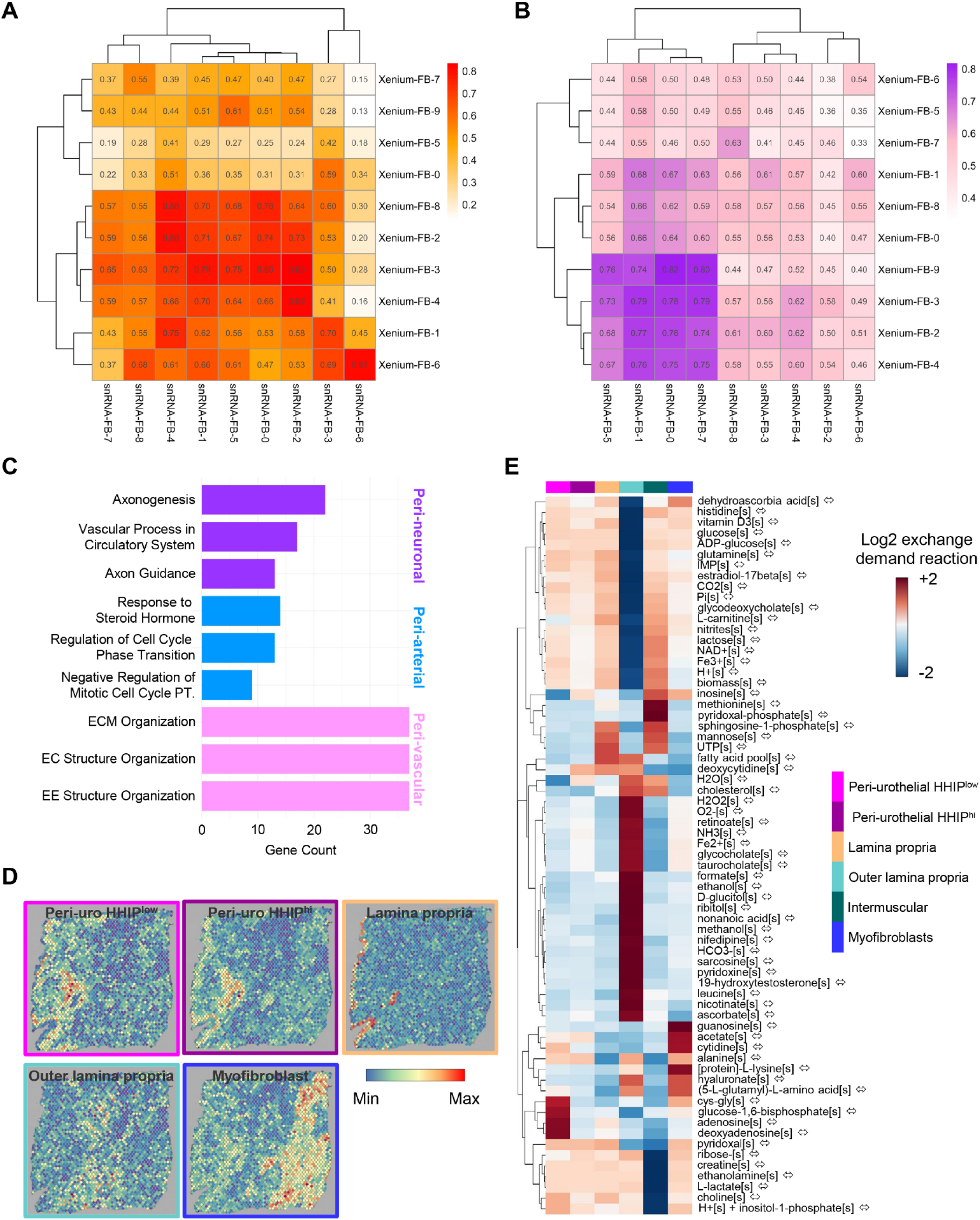
Cross-platform mapping and characterization of bladder fibroblast subpopulations. (**A-B**) Heatmaps showing correlation between fibroblast subpopulations identified in Xenium (xFBs) and multimodal snRNA-seq clustering (snRNA-FBs), based on shared gene (**A**) and poly(A) isoform (**B**) expression profiles from the Xenium panel and snRNA-seq normalized RNA assay. (**C**) Bar plot showing enriched gene ontology (GO) terms for peri-neuronal, peri-arterial, and peri-vascular fibroblasts, based on differentially expressed genes from snRNA-seq. (**D**) Visium spatial transcriptomics of an independent bladder tissue sample demonstrating reproducibility of the fibroblast distribution pattern using only metabolism-related genes curated from the Recon3D database. (**E**) Heatmap showing relative enrichment of exchange demand reactions for nutrient metabolism across fibroblast layers, computed using METAFlux. Log_2_-transformed exchange demand values are plotted. Positive values indicate nutrient uptake, and negative values indication release.

**Table S1. (separate file)**

Sample and sequencing statistics with cell counts

**Table S2. (separate file)**

snRNA-seq cell cluster markers

**Table S3. (separate file)**

Harmonized poly(A) peak reference

**Table S4. (separate file)**

Alternative polyadenylation events between bladder cell types

**Table S5. (separate file)**

Xenium cell QC statistics

**Table S6. (separate file)**

Xenium cell cluster markers

**Table S7. (separate file)**

Cell neighborhood analysis using Xenium data

**Table S8. (separate file)**

Fibroblast cell cluster markers from multimodal clustering using gene and isoform information

**Table S9. (separate file)**

Multimodal fibroblast cluster GO biological processes analysis results

